# Degradation of Polycomb Repressive Complex 2 with an EED-targeted Bivalent Chemical Degrader

**DOI:** 10.1101/676965

**Authors:** Frances Potjewyd, Anne-Marie W. Turner, Joshua Beri, Justin M. Rectenwald, Jacqueline L. Norris-Drouin, Stephanie H. Cholensky, David M. Margolis, Kenneth H. Pearce, Laura E. Herring, Lindsey I. James

## Abstract

Protein degradation via the use of bivalent chemical degraders provides an alternative strategy to block protein function and assess the biological roles of putative drug targets. This approach capitalizes on the advantages of small molecule inhibitors while moving beyond the restrictions of traditional pharmacology. Herein we report a first-in-class chemical degrader (UNC6852) that targets Polycomb Repressive Complex 2 (PRC2). UNC6852 contains an EED226 derived ligand and a ligand for VHL which bind to the WD40 aromatic cage of EED and CRL2^VHL^, respectively, to induce proteasomal degradation of PRC2 components, EED, EZH2, and SUZ12. Degradation of PRC2 with UNC6852 blocks the histone methyltransferase activity of EZH2, decreasing H3K27me3 levels in HeLa cells and diffuse large B-cell lymphoma (DLBCL) cells containing an EZH2^Y641N^ gain-of-function mutation. UNC6852 degrades both wild type EZH2 and EZH2^Y641N^, and additionally displays anti-proliferative effects in this cancer model system.

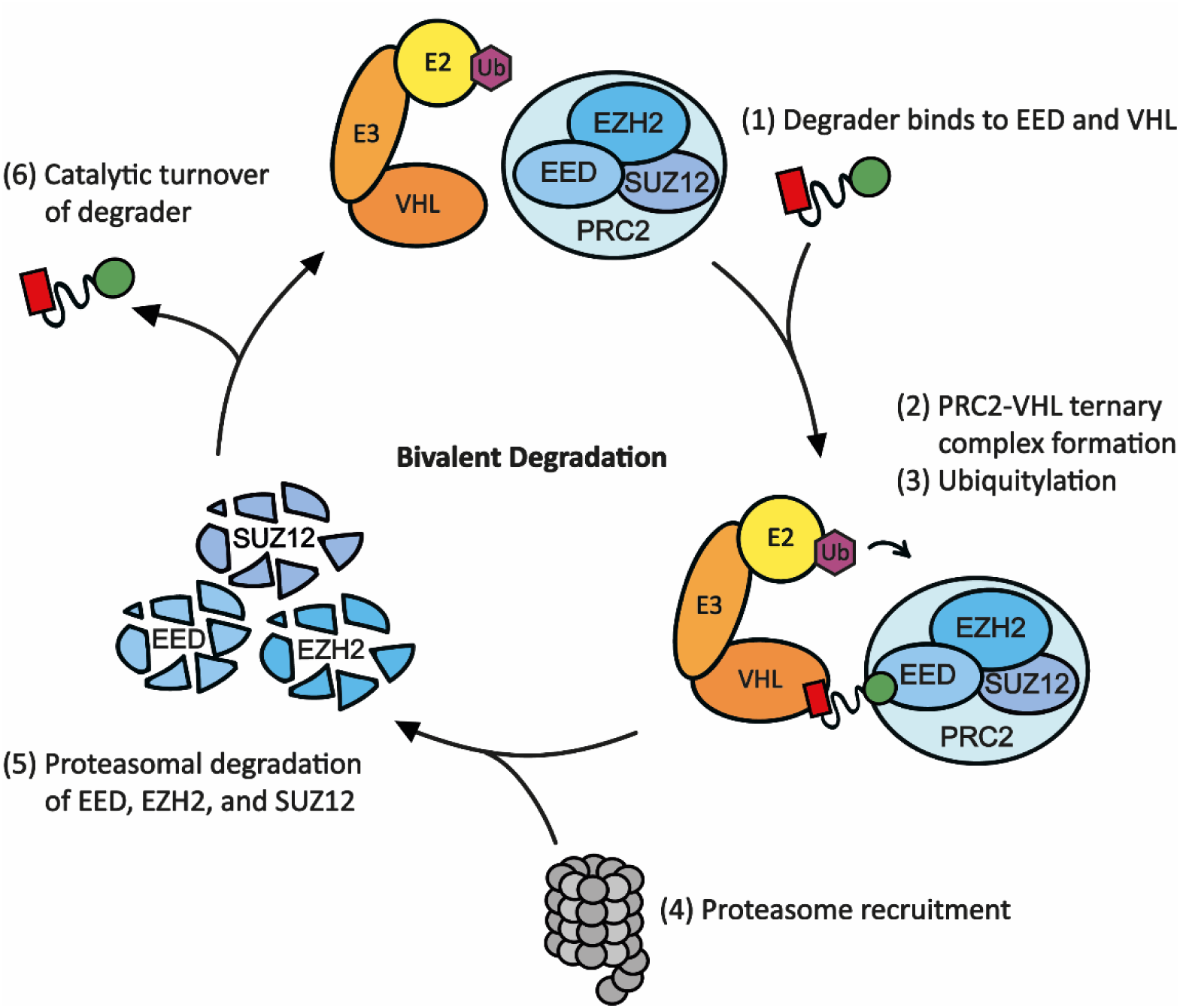

## INTRODUCTION

Polycomb Repressive Complex 2 (PRC2) is a multicomponent complex with histone methyltransferase (HMT) activity that installs and maintains mono-through trimethylation at histone 3 lysine 27 (H3K27). H3K27 trimethylation (H3K27me3) is a key mechanism responsible for gene repression (Ferrari *et al*., 2014). The catalytic activity of PRC2 is dependent on the formation of a complex containing three core subunits: Embryonic Ectoderm Development (EED), Enhancer of Zeste Homolog 1 (EZH1) or EZH2, and Suppressor of Zeste Homolog 12 (SUZ12) (Margueron and Reinberg, 2011). EZH1 and EZH2 share significant sequence homology and both HMTs can be incorporated into PRC2 to generate an active complex; however, EZH1 has a lower abundance and often lesser HMT activity as compared to EZH2 (Margueron *et al*., 2008; Lee *et al*., 2018). Other proteins commonly associated with PRC2 include Jumonji and AT-rich interacting Domain 2 (JARID2), (Peng *et al*., 2009) PHD finger protein 19 (PHF19), and AE Binding Protein 2 (AEBP2) (Hyun *et al*., 2017). Structural elucidation of PRC2 revealed an intricate network of protein-protein interactions between EED, EZH2 and SUZ12 which are necessary for PRC2 catalytic activity (Liu, 2015; Justin *et al*., 2016; Kasinath *et al*., 2018; Poepsel, Kasinath and Nogales, 2018). Specifically, EED recognition of H3K27me3 via its WD40 domain serves to stabilize the stimulation responsive motif (SRM) of EZH2 and allosterically activates the SET domain of EZH2 for trimethylation of H3K27 on adjacent nucleosomes (Justin *et al*., 2016).

PRC2 has been reported as both an oncogene and suppressor of tumorigenesis in an assortment of cancer types (Gan *et al*., 2018). EZH2, EED, and SUZ12 are commonly upregulated in certain cancers such as breast, colorectal, and prostate cancer (Liu *et al*., 2015; Gan *et al*., 2018). Overexpression of EZH2 and elevated levels of H3K27me3 have been linked to both increased cell proliferation and chemotherapy resistance which can result in low survival rates. EED, EZH2, and SUZ12 are also susceptible to mutations in cancer. For example, EZH2 gain-of-function mutations are commonly associated with lymphomas. Heterozygous EZH2 gain-of-function mutations in the C-terminal SET domain occur at Y641, A677, and A687, and lead to EZH2 hyperactivity, an increase in global H3K27me3 levels, and aberrant gene repression (Veneti, Gkouskou and Eliopoulos, 2017). Diffuse Large B-cell Lymphomas (DLBCL) commonly harbour these mutations, marking EZH2 as an important target for therapeutic intervention (McCabe *et al*., 2012).

Effective inhibition of PRC2 catalytic activity has been achieved by targeting both EED and EZH2. While initial efforts were focused on developing inhibitors of the catalytic SET domain of EZH2 (Genta, Pirosa and Stathis, 2019), it was recently demonstrated that small molecule antagonists of the EED WD40 domain could phenocopy EZH2 inhibitors due to the critical role of EED in regulating PRC2 activity (He *et al*., 2017; Qi *et al*., 2017). EED and EZH2 inhibition have each been shown to reduce global H3K27me3 levels and result in antiproliferative effects in EED and EZH2 wild type (WT) cancer cell lines, as well as cell lines with EZH2 gain-of-function mutations (Xu *et al*., 2015; He *et al*., 2017; Shortt *et al*., 2017; Lee *et al*., 2018). EZH2 inhibitors that bind the SET domain include chemical probes such as UNC1999, as well as several compounds in clinical development including GSK126, EPZ-6438 (Tazemostat), CPI-1205, and DS-3201b (Valemostat), which have been particularly effective in lymphomas with activating EZH2 mutations (McCabe *et al*., 2012; Konze *et al*., 2013; Dilworth and Barsyte-Lovejoy, 2019; Genta, Pirosa and Stathis, 2019). More recently, EED chemical probes EED226 and A-395 were reported, and currently MAK683, an analogue of EED226, is in the clinic for similar applications (He *et al*., 2017; Huang *et al*., 2017; Dilworth and Barsyte-Lovejoy, 2019). Resistance to EZH2 inhibitors has been observed in the clinic and is one limitation to this class of SAM-competitive molecules; however, EED antagonists have the potential to overcome this acquired resistance (Brooun *et al*., 2016; Lee *et al*., 2018). Overall, targeting PRC2 for cancer treatment has been shown to be an effective strategy, yet new approaches are needed to overcome observed resistance to EZH2 inhibitors and to develop novel therapeutics.

Bivalent chemical protein degraders, otherwise known as PROTACs™, are molecules designed to degrade a specific endogenous protein of interest (POI) by harnessing the E3 Ubiquitin ligase pathway (Cromm and Crews, 2017; Salami and Crews, 2017). Bivalent protein degraders are composed of a ligand for the desired POI, an E3 ligase ligand, and an optimized linker connecting the two ligands. The most extensively used E3 ligase recruiting ligands include VH032 and pomalidomide, which are responsible for recruitment of von-Hippel Lindau (VHL) as part of the CRL2^VHL^ E3 ubiquitin ligase complex and cereblon (CRBN) as part of the CRL4^CRBN^ E3 ubiquitin ligase complex, respectively (Fischer *et al*., 2014; Cardote, Gadd and Ciulli, 2017; Cromm and Crews, 2017). The linker region typically consists of a flexible alkyl or polyethylene glycol (PEG) moiety, although other linkers have been explored. Linked ligands bring the POI into close proximity with the E3 ligase recruiting protein to form a ternary complex, which allows the E3-ligase Cullin ring complex to ubiquitylate a lysine residue on the POI, thereby tagging the protein for proteasomal degradation (Gadd *et al*., 2017). Positive cooperativity of ternary complex formation between these proteins and the subsequent ubiquitylation of an available lysine are both important factors for efficient proteasomal degradation. Additionally, chemical degraders act catalytically which compensates for their inherently low cell permeability (Bondeson *et al*., 2015; Riching *et al*., 2018). Because they are catalytic and don’t require very high-affinity for their POI, bivalent chemical degraders have the potential to facilitate degradation of previously ‘undruggable’ targets and represent a promising therapeutic strategy. Just recently, the first PROTAC™ entered the clinic for the treatment of metastatic castration-resistant prostate cancer (Mullard, 2019), demonstrating that the anticipated pharmacokinetic challenges due to their high molecular mass can be overcome. Due to the availability of ligands for both EZH2 and EED, we postulated that the development of bivalent degraders could be an effective alternative strategy to inhibit PRC2 function.

Herein we describe the design, synthesis, and evaluation of a novel PRC2 bivalent chemical degrader based on the potent EED ligand EED226. To the best of our knowledge, our efforts have yielded the discovery of the first PRC2 degrader, which effectively degrades EED, EZH2, and SUZ12 in a VHL-dependent fashion, reduces H3K27me3 levels, and decreases proliferation of DB cells, a DLBCL cell line harboring the EZH2^Y641N^ mutant. Together, these results demonstrate the feasibility of developing PRC2-targeted degraders to block PRC2 function, to interrogate PRC2 biology, and as potential therapeutics.

## RESULTS

### Design and Synthesis of EED-Targeted Bivalent Degraders

Based upon the successful development of potent ligands for EED which function as allosteric inhibitors of PRC2 and the emerging field of bivalent chemical degraders, we designed and synthesized a series of heterobifunctional EED-targeted chemical degraders. These compounds are comprised of an analogue of a known EED ligand, EED226, and VH032-amine, a ligand which has been successfully employed in numerous examples for CRL2^VHL^ recruitment (Figure 1) (Frost *et al*., 2016; Qi *et al*., 2017; An and Fu, 2018; Zou, Ma and Wang, 2019). We first needed to identify an exit vector on EED226 that would be synthetically amenable to functionalization with a linker moiety without a significant loss in potency. A crystal structure of EED226 bound to the WD40 domain of EED indicated that the sulfone moiety of EED226 is solvent exposed, providing a potential site for functionalization (PDB: 5GSA, Qi *et al*. 2017). VHL ligands have several known functionalization sites based on their prior incorporation into bivalent degraders providing multiple possible exit vectors. Importantly, the exit vector chosen can have a large impact on ternary complex formation (Cromm and Crews, 2017; Chan *et al*., 2018; Smith *et al*., 2019). Functionalization off of the terminal amine of VH032 has been extensively used in the design of bivalent degraders so we chose this position for linker appendage (Frost *et al*., 2016; Chan *et al*., 2018; Girardini *et al*., 2019). To connect the two ligands, different length alkyl (UNC6851-UNC6853) and PEG linkers (UNC6845-UNC6847) were incorporated to assess the distance required to induce successful EED degradation upon formation of the EED-degrader-VHL ternary complex (Table 1) (Cyrus *et al*., 2011). To enable this approach, we synthesized a carboxylic acid functionalized EED ligand via a Suzuki-Miyaura reaction with (4-(methoxycarbonyl)phenyl)boronic acid and 8-bromo-*N*-(furan-2-ylmethyl)-[1,2,4]triazolo[4,3-*c*]pyrimidin-5-amine (**1**) and subsequent basic hydrolysis to yield (**2**) (Scheme S1). VH032-amine (**3**) was reacted with the various *N*-Boc alkyl and PEG linkers followed by deprotection (**4** – **9**). Assembly of the final compounds was achieved by an amidation reaction to afford UNC6851, UNC6852, UNC6853, UNC6845, UNC6846, and UNC6847 (Scheme S2).

**Table 1.**
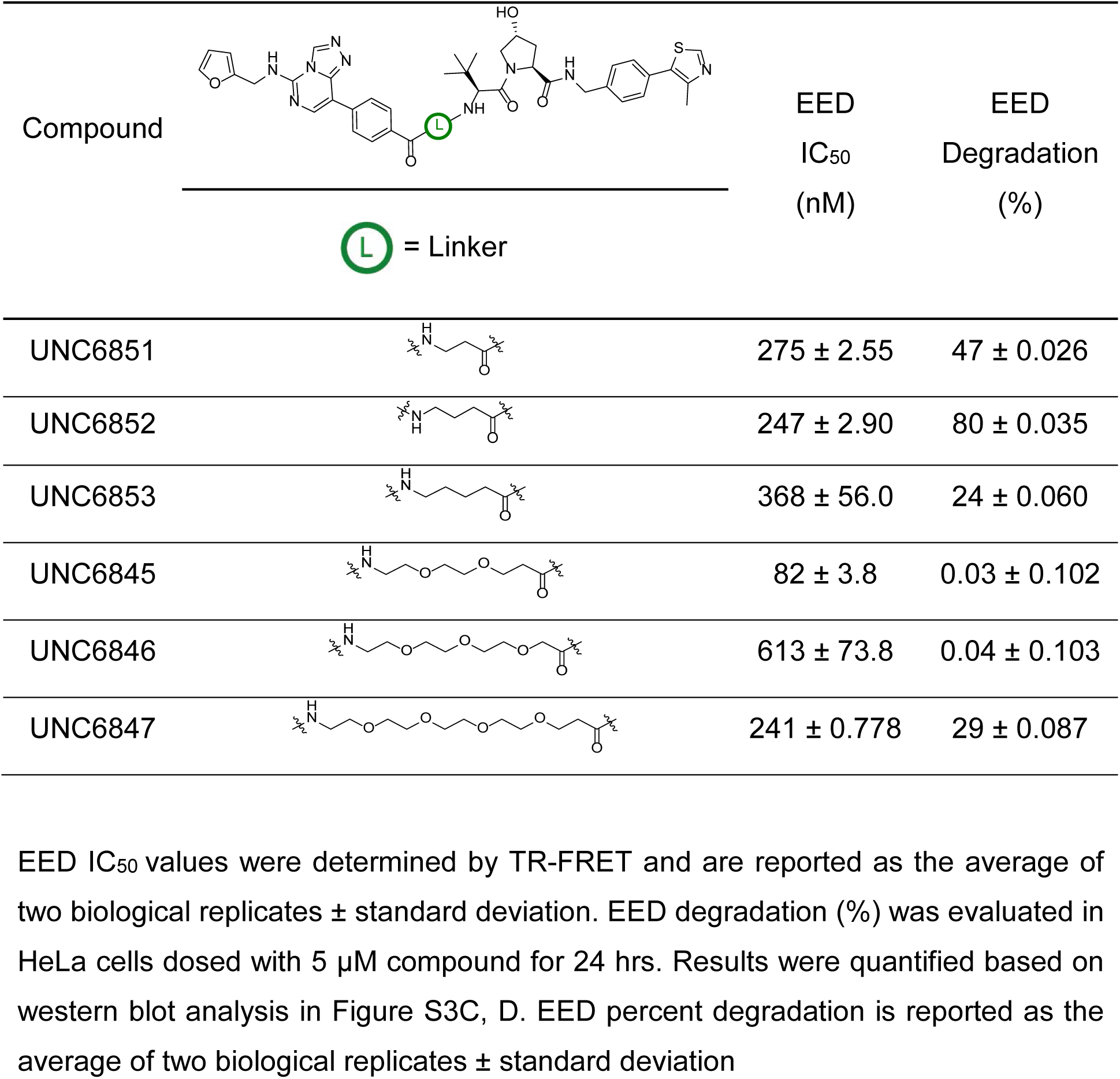
Analysis of the extent of EED binding and degradation with 6 EED-targeted degraders.

| Compound |  | EED<br>IC<sub>50</sub><br>(nM) | EED<br>Degradation<br>(%) |
| --- | --- | --- | --- |
| UNC6851 |  | 275 ± 2.55 | 47 ± 0.026 |
| UNC6852 |  | 247 ± 2.90 | 80 ± 0.035 |
| UNC6853 |  | 368 ± 56.0 | 24 ± 0.060 |
| UNC6845 |  | 82 ± 3.8 | 0.03 ± 0.102 |
| UNC6846 |  | 613 ± 73.8 | 0.04 ± 0.103 |
| UNC6847 |  | 241 ± 0.778 | 29 ± 0.087 |

**Figure 1.**
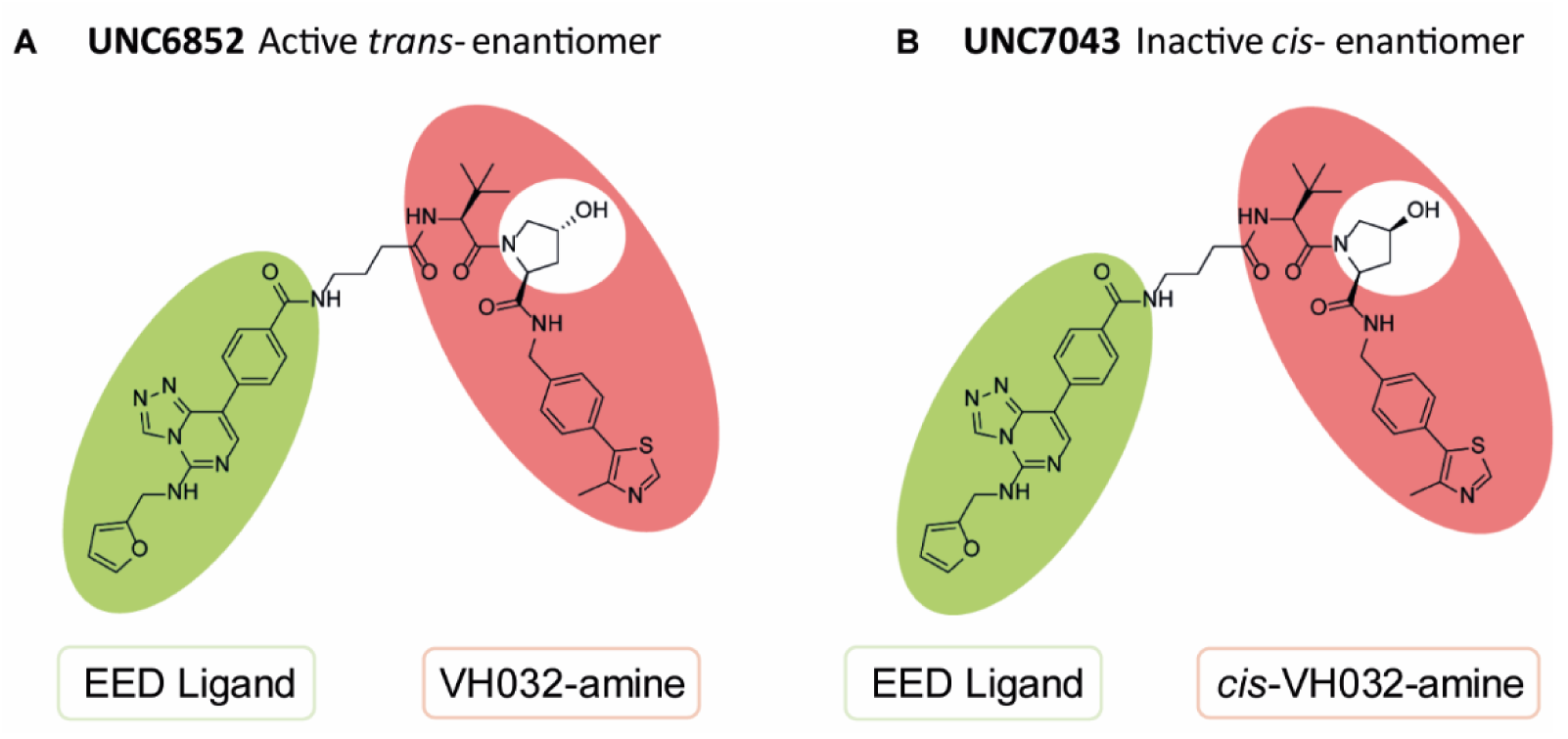
Chemical structures of UNC6852 and UNC7043. (**A**) UNC6852 is a bivalent chemical degrader of PRC2 containing an EED ligand (green) and a VHL ligand (coral). (**B**) UNC7043 is a corresponding inactive control compound which contains a *cis*-hydroxyproline amino acid, abrogating binding to VHL.

Initially we confirmed that our bivalent molecules still potently bound to the WD40 domain of EED via a time-resolved fluorescence resonance energy transfer (TR-FRET) assay. In this assay we used 6XHis-tagged recombinant EED (residues 1 – 441) and a biotinylated EED ligand previously developed in our lab (UNC5114-biotin, Barnash *et al*. 2017), with a fluorophore labelled anti-6XHis antibody (acceptor) and europium labelled streptavidin (donor), respectively. EED226 was used as a positive control and displayed potencies comparable to literature reported values (Figure S1, IC_50_ = 45 nM; reported IC_50_ = 22 nM) (Huang *et al*., 2017). We also synthesized a negative control EED ligand, UNC5679, which showed no significant binding within the concentrations tested and is > 200-fold less potent than EED226 (Figure S1, IC_50_ = >10 μM; reported IC_50_ = 20.49 μM) (Huang *et al*., 2017). Alkyl linked compounds UNC6851, UNC6852, and UNC6853 showed a 6-fold, 5.5-fold, and 8-fold loss in potency compared to EED226, respectively. PEG linked compounds UNC6845, UNC6846, and UNC6847 revealed a 2-fold, 14-fold, and 5-fold loss in potency, respectively. Overall, these data confirm that our bivalent molecules are sufficiently potent binders of the WD40 domain of EED, and therefore should be able to engage EED as the first step in initiating the E3-ligase mediated proteasomal degradation pathway.

### UNC6852 Mediates PRC2 Degradation

Next, we sought to assess the ability of our six bivalent molecules to enable EED degradation. To do so, we first performed extensive antibody validation studies utilizing overexpression systems to identify EED and EZH2 antibodies that were both compatible with the Jess™ system for automated protein analysis (ProteinSimple) and suitable for follow-up studies (Figure S2). HeLa lysates were then generated from cells treated with bivalent degraders (5 μM) for 4, 24, and 48 hrs and screened on the Jess™ system (Figure S3A, B), which allows for the analysis of protein degradation in a more high-throughput fashion than traditional western blotting experiments. Due to the close proximity of EZH2 residues to the EED226 binding site, we speculated that EED226-derived degraders may additionally facilitate EZH2 degradation, and therefore both EED and EZH2 protein levels were monitored. Encouragingly, these data indicated that UNC6851 and UNC6852 resulted in a decrease in the levels of both EED and EZH2 at 24 hrs, with UNC6852 having a more pronounced effect than UNC6851 at shorter and longer time points (4 and 48 hours, Figure S3A, B). These compounds differ by a single CH_2_ group in the linker, with UNC6851 containing a 2-methylene linker and UNC6852 a 3-methylene linker. In contrast, significant degradation was not observed with UNC6853 which contains a slightly longer 4-methylene linker, highlighting that even minor variations in a linker moiety can significantly impact degradation efficiency. The bivalent molecules with PEG linkers (UNC6845, UNC6846, and UNC6847), all of which are longer than the 4-methylene linker of UNC6853, were similarly unable to alter the levels of EED or EZH2 under these conditions. To validate these results, we performed traditional western blot analysis, evaluating EED and EZH2 protein levels after treatment with each degrader for 24 hours. UNC6852 was again identified as the most proficient degrader of EED (80% degradation, Table 1) and EZH2 (76%, Figure S3C, D) under these conditions.

To further investigate the degradation potential of UNC6852, we evaluated EED and EZH2 levels upon treatment with UNC6852 in a dose response format at 24 hrs and over various times at a fixed concentration (10 μM) by western blot analysis (Figure 2, Figure S4). Upon treatment of HeLa cells with UNC6852, no cellular toxicity was observed at concentrations up to 30 μM. UNC6852 was capable of degrading EED and EZH2 to varying extents at different concentrations and time points. EED and EZH2 degradation occurred at similar concentrations of UNC6852, with DC_50_ values (the concentration at which 50% degradation was observed) of 0.79 ± 0.14 μM and 0.3 ± 0.19 μM, respectively (Figures S4B, G). The maximal degradation observed (D_max_) was slightly higher for EED (92%) than EZH2 (75%) and interestingly, EED was also degraded at earlier time points than EZH2, with an apparent half-life (t_1/2_) of 0.81 ± 0.30 hours and 1.92 ± 0.96 hours, respectively (Figures S4A, G).

**Figure 2.**
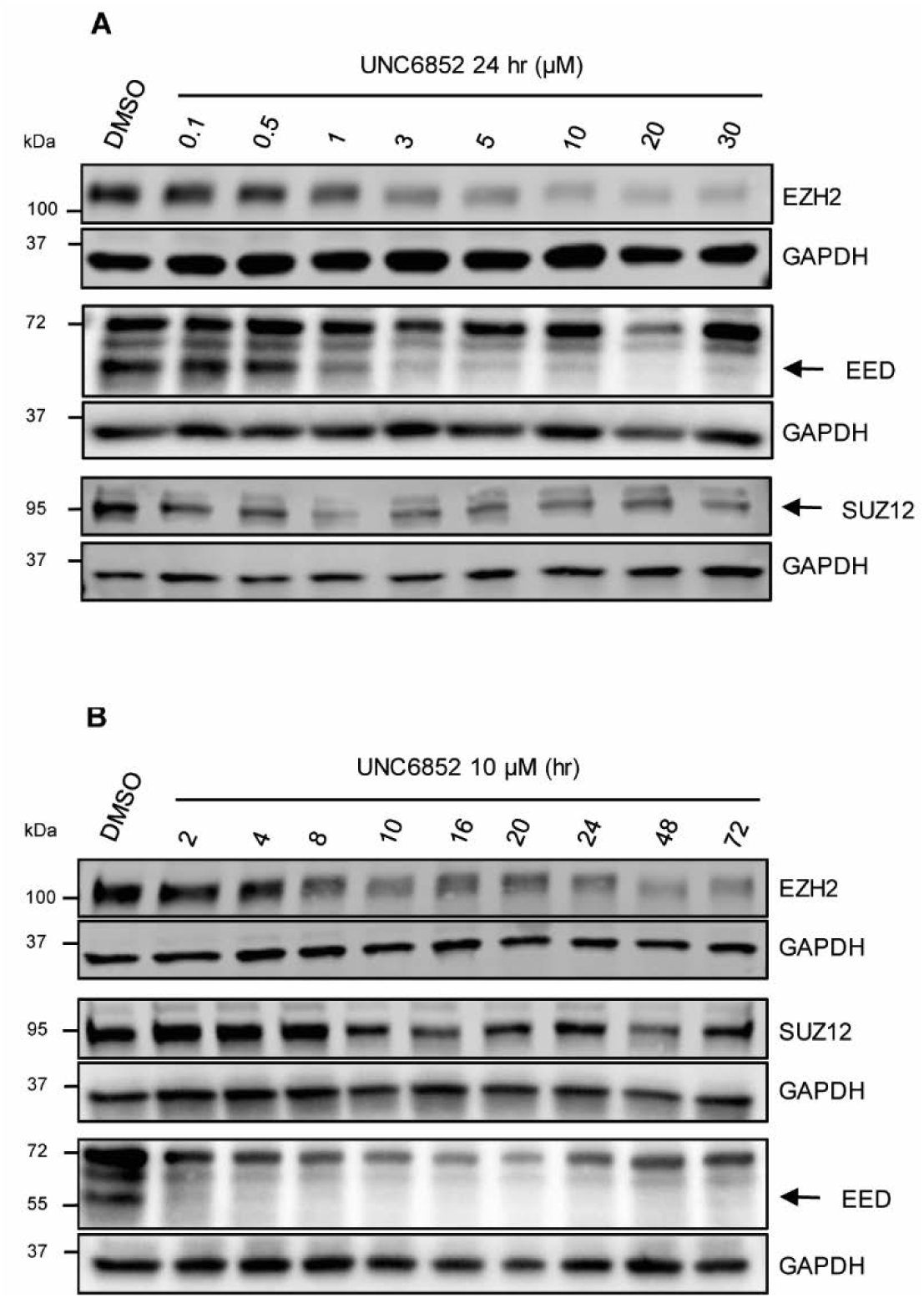
UNC6852 degrades PRC2 components EED, EZH2 and SUZ12 in HeLa cells. (**A**) Western blot analysis of PRC2 components following UNC6852 treatment in a dose response fashion (0 – 30 μM, 24 hours). (**B**) Western blot analysis of PRC2 components following treatment of UNC6852 (10 μM) from 2 to 72 hrs. Data is representative of at least two biological replicates. Quantification of these results are reported in Supplementary Figure S4.

SUZ12 is the third core component of PRC2, and thus we were equally interested in determining if UNC6852 can effectively degrade SUZ12. In the dose response and time course studies described above, we found that, SUZ12 was degraded to a lesser extent than both EED and EZH2 by UNC6852 (Figure 2, Figure S4). We were unable to calculate the DC_50_ and half-life for SUZ12 due to a maximal degradation of only 22%.

### UNC6852 Facilitates PRC2 Degradation via VHL Recruitment

To confirm that UNC6852 is inducing degradation of PRC2 via the ubiquitin-proteasomal degradation pathway induced by CRL2^VHL^ E3-ligase recruitment, we utilized proteasome inhibitors and an inactive heterobifunctional control compound (UNC7043, Figure 1). We designed and synthesized UNC7043, which is structurally identical to UNC6852 except for the fact that it contains the opposite enantiomer at the hydroxyproline moiety on the VHL ligand (Scheme S2). This subtle change to VH032 disables ligand binding to VHL, and hence when incorporated into a bivalent molecule, no longer recruits VHL. As expected, UNC7043 treatment did not degrade EED, EZH2, or SUZ12 when HeLa cells were dosed at 10 μM for 24 hrs (Figure 3A). This result further established that UNC6852-mediated degradation of PRC2 is occurring via the CRL2^VHL^ based ubiquitin-proteasomal degradation pathway.

**Figure 3.**
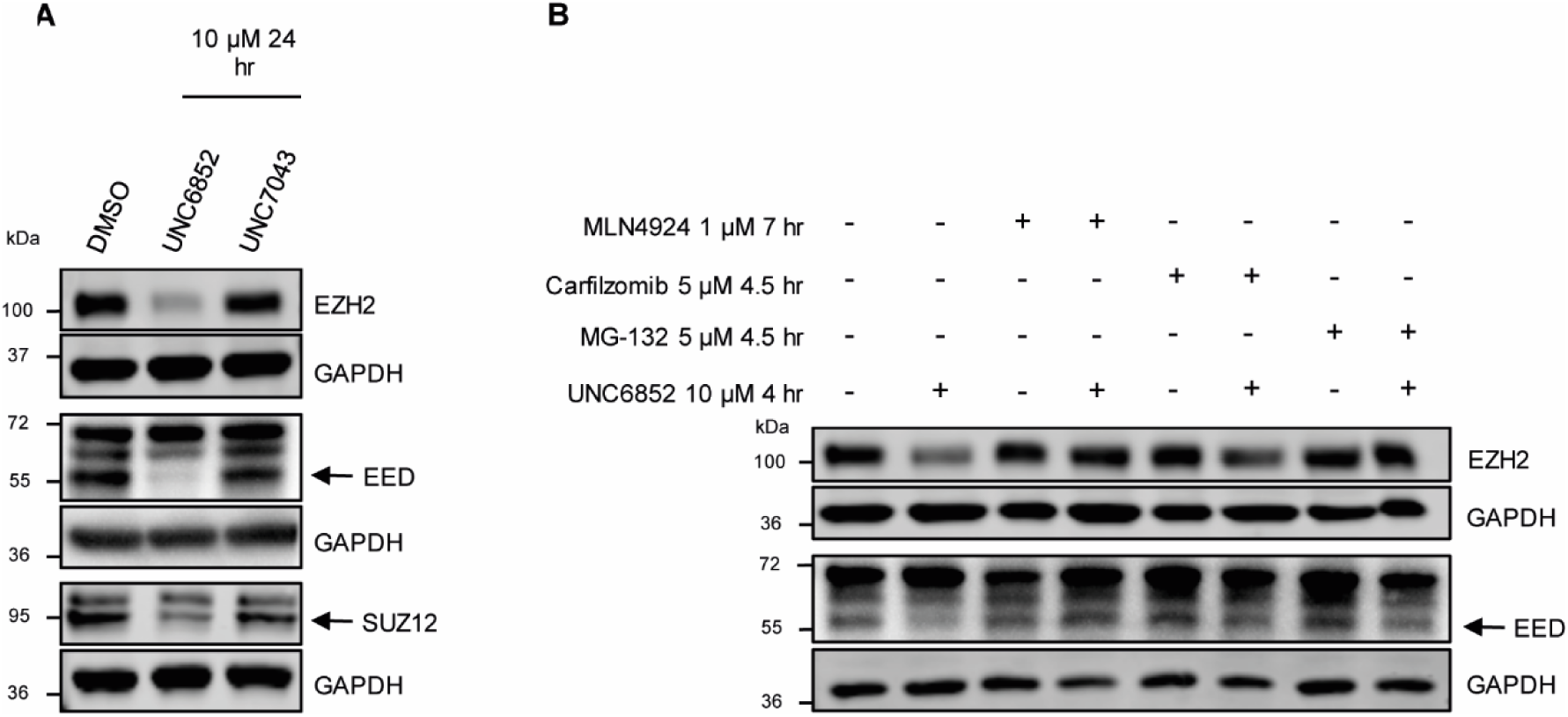
PRC2 components are not degraded upon treatment with proteasome inhibitors or negative control compound UNC7043. (**A**) Western blot analysis of PRC2 components upon treatment of HeLa cells with UNC6852 and negative control compound UNC7043 (10 μM for 24 hours). (**B**) Western blot analysis of PRC2 components in HeLa cells pre-treated with proteasome inhibitors MLN4924 (1 μM for 7 hours), Carfilzomib and MG-132 (5 μM for 4.5 hours), followed by UNC6852 4 hrs at 10 μM. Data is representative of at least two biological replicates.

Additionally, pre-treatment with proteasome inhibitors MLN4924, Carfilzomib, and MG-132 prior to addition of UNC6852 effectively blocked EED and EZH2 degradation, again confirming the proposed degradation mechanism (Figure 3B). Specifically, HeLa cells were pre-treated for 7 hrs with MLN4924 (Pevonedistat) and 4.5 hrs with Carfilzomib or MG-132 to halt cellular ubiquitylation mechanisms prior to addition of UNC6852 for 4 hrs. Degradation effects could not be evaluated at longer time points due to the toxicity inherent to these proteasome inhibitors (Maniaci *et al*., 2017; Huang *et al*., 2018). While it has been previously reported that proteasome inhibitor treatment can decrease endogenous EZH2 levels, treatment of HeLa cells with proteasome inhibitors alone did not change EZH2 levels under these conditions (Rizq *et al*., 2017).

### UNC6852 selectively degrades EED and EZH2

To assess the effects of UNC6852 treatment on cellular protein levels more broadly, we performed global proteomics experiments using tandem mass tag (TMT) quantification comparing HeLa cells treated with UN6852 (10 μM, 24 hrs) to DMSO treated control cells. Whole proteome analysis resulted in the identification of >60,000 peptides corresponding to 5,452 quantifiable proteins. Notably, these data revealed that EED and EZH2 were selectively degraded by UNC6852 within the proteome (Figure 4). Significant degradation was defined by a p-value of <0.01 and a log2 fold change ratio of −0.5 (UNC6852 treated/DMSO treated). Although SUZ12 did not meet these criteria (log2 fold change = −0.34), modest SUZ12 degradation (21%) was observed which is consistent with our previously determined D_max_ value via western blot analysis (D_max_ = 22%, Figure S4). Importantly, in parallel studies with Hela cells treated with negative control degrader, UNC7043 (10 μM, 24 hrs), EED, EZH2, and SUZ12 were not similarly degraded, confirming that selective PRC2 degradation is occurring via the E3-ligase ubiquitylation pathway.

**Figure 4.**
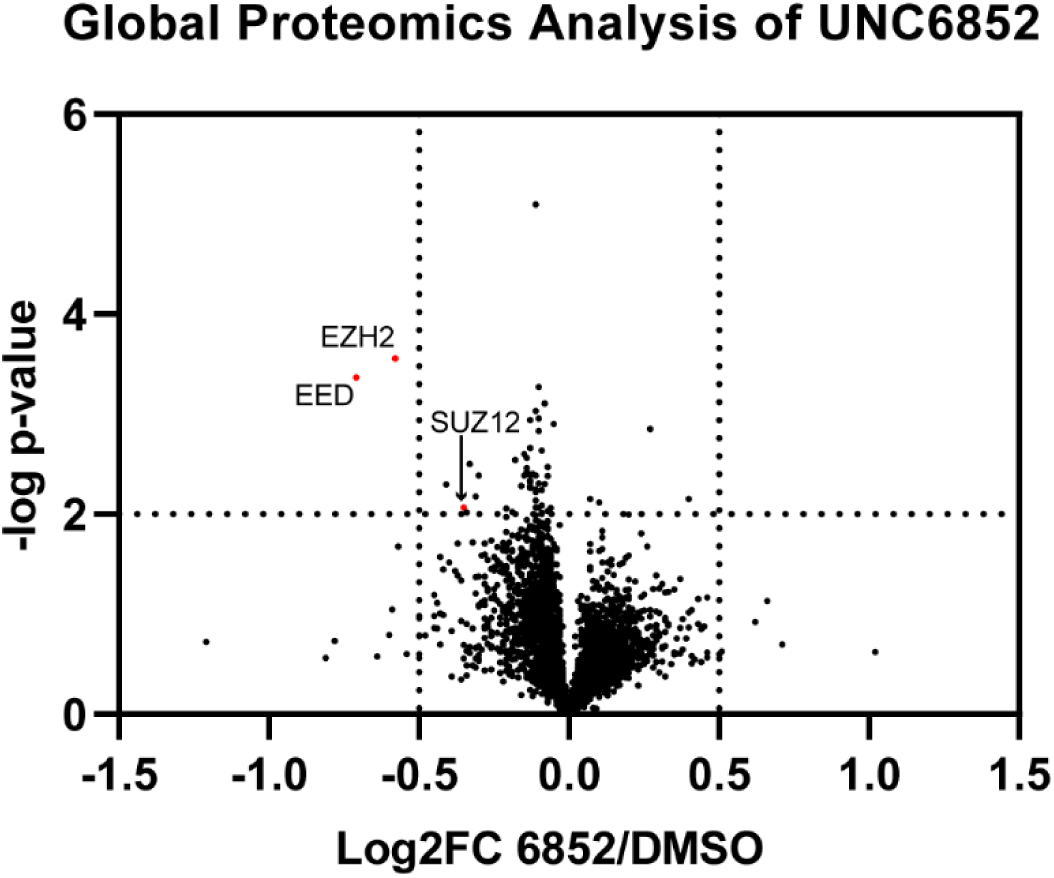
UNC6852 selectively degrades PRC2. Quantitative proteomics results showing relative abundance of proteins in HeLa cells treated with DMSO, UNC6852, or UNC7043 (10 μM, 24 hrs). Of the total 5,452 quantifiable proteins, EED and EZH2 were selectively degraded by UNC6852 within the proteome. Significant degradation was defined by a p-value of <0.01 and a log2 fold change ratio of −0.5 (UNC6852 treated/DMSO treated). Data shown are three replicates measured in a single 10-plex TMT experiment.

### UNC6852 reduces H3K27me3 levels and DLBCL cell proliferation

We next sought to investigate the effects of PRC2 degradation on H3K27me3 levels and cellular proliferation. We first treated HeLa cells with UNC6852 and EED226 over a time course of 24 to 72 hrs at 10 μM to monitor H3K27me3 levels by western blot. As expected, UNC6852 resulted in a decrease in protein levels of both EED and EZH2 over these time points, whereas EED226 had no effect (Figure S5). Importantly, UNC6852 and EED226 led to a comparable decrease in H3K27me3 levels, with H3K27me3 reduced by 51% and 53%, respectively, after 72 hours (Figure S5B).

Next, we were interested in evaluating the sensitivity of diffuse large B-cell lymphoma (DLBCL) cell lines that contain a heterozygous EZH2^Y641N^ gain-of-function mutation to UNC6852. The EZH2^Y641N^ mutation leads to an increase in H3K27me3 levels due to PRC2 hyperactivity (Xu *et al*., 2015). First, we investigated the effect of UNC6852 on PRC2 degradation in DB cells in a dose-dependent manner at 24 hours (Figure 5A). We observed degradation of EED and EZH2/EZH2^Y641N^ in DB cells with a similar half maximal degradation concentration as in HeLa cells (DC_50_ = 0.61 ± 0.18 μM and 0.67 ± 0.24 μM, respectively). In contrast to prior results in HeLa cells, where we observed partial degradation, EZH2/EZH2^Y641N^ and EED were both completely degraded by UNC6852 (D_max_ = 96% and 94%, respectively). Additionally, SUZ12 was also degraded to a much larger extent in DB cells. The maximal degradation of SUZ12 was 3.7-fold higher (D_max_ = 82%) than in HeLa cells, with a calculated half maximal degradation concentration of 0.59 ± 0.17 μM. As expected, treatment with UNC7043 in DB cells did not affect levels of these proteins (Figure 5B). Degradation of PRC2 by UNC6852 in DB cells also significantly reduced H3K27me3 levels, with a 71% loss of H3K27me3 after 72 hours (Figure 5C, D). Overall, UNC6852 potently degrades the core components of PRC2 and results in a concomitant loss of H3K27me3 in DB cells with an EZH2 gain-of-function mutation.

**Figure 5.**
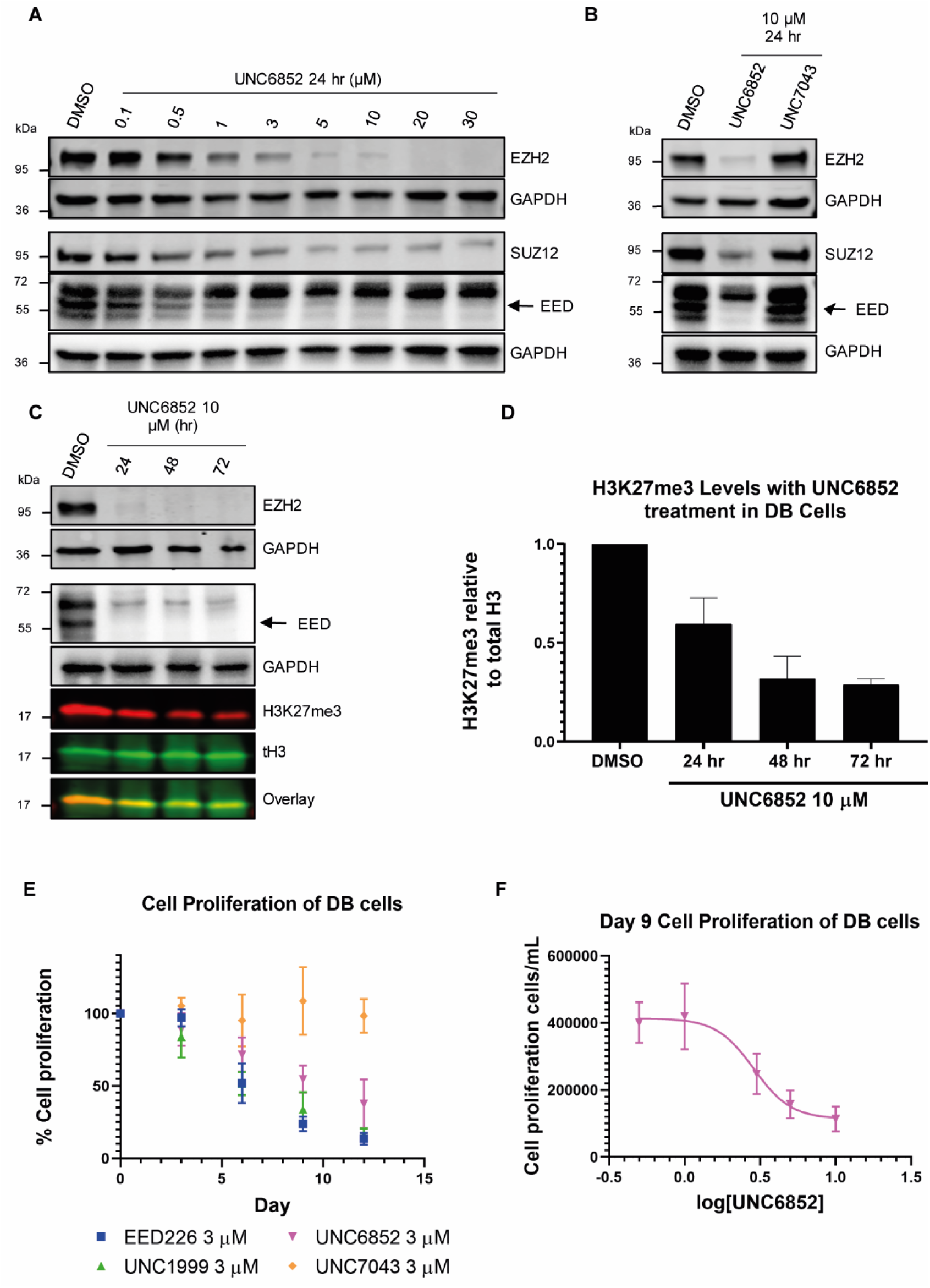
UNC6852 degrades PRC2, reduces H3K27me3 levels, and decreases proliferation in an EZH2^Y641N^ DB cell line. (**A**) Western blot analysis of the degradation of EED, EZH2, and SUZ12 in DB cells containing a heterozygous EZH2^Y641N^ mutation treated with UNC6852 (0.1 – 30 μM for 24 hours). (**B**) Western blot analysis following treatment of DB cells with UNC6852 or negative control compound UNC7043 (10 μM for 24 hours). (**C**) Western blot analysis of PRC2 components and H3K27me3 in DB cells treated with UNC6852 in a time dependent fashion (10 μM for 24, 48 and 72 hours). (**D**) Quantification of H3K27me3 levels relative to total H3 in **C**. DMSO control was normalized to 1. (**E**) DB cell line proliferation data upon treatment with EED226, UNC1999, UNC6852, and UNC7043 (3 μM) reported relative to DMSO treatment. Corresponding cell viability data is shown in Supplementary Figure S6A. (**F**) UNC6852 displays a concentration dependent inhibition of DB cell proliferation after 9 days of treatment (0.5 – 10 μM). Corresponding cell viability data is shown in Supplementary Figure S6B.

Furthermore, when DB cells were treated with UNC6852 for 12 days a robust anti-proliferative effect was observed (Figure 5E). Cells were similarly treated with EED226 or UNC1999, a potent EZH1/2 inhibitor, both of which have been shown to effectively reduce DLBCL cell proliferation (Konze *et al*., 2013; Huang *et al*., 2017). After 9 days, UNC6852 displayed a concentration dependent inhibition of cell proliferation, similar to EED226 and UNC1999, with an EC_50_ of 3.4 ± 0.77 μM (Figure 5F). Additionally, overall cell toxicity was significantly less with UNC6852 (95% viable cells) as compared to EED226 (67% viable cells) and UNC1999 (67% viable cells) after 12 days (Figure S6A). Interestingly, the negative control compound UNC7043 which is unable to bind VHL had no effect on cell proliferation despite containing a potent EED ligand (Figure 5E). This result reaffirms that the difference in proliferative effects between EED226 and UNC7043 in DB cells is likely due to the lack of permeability inherent to most bivalent degraders and that catalytic degradation is required for efficacy of UNC6852. Concordantly, it can be concluded that the anti-proliferative effects seen with UNC6852 are due to PRC2 degradation as opposed to EED inhibition alone.

## DISCUSSION

Herein we report the first example of PRC2 degradation based on our discovery of an EED-targeted bivalent chemical degrader. We show that UNC6852 potently binds EED *in vitro*, degrades EED and other PRC2 components in a highly selective fashion, inhibits PRC2 catalytic activity resulting in decreased H3K27me3 levels, and has antiproliferative effects in DB cell lines. We demonstrate that UNC6852 effectively degrades PRC2 components EED, EZH2, and SUZ12 via VHL recruitment and the E3-ligase proteasome degradation pathway.

To achieve efficient degradation, productive ternary complex formation with EED and VHL, as well as subsequent ubiquitylation of an available lysine residue, are essential. In common with other chemical degraders, we found that the linker incorporated to bridge the EED and VHL ligands was critical. UNC6852 contains a short alkyl linker of only three methylene groups, and we were surprised to find that the addition of a fourth methylene group (UNC6853) was sufficient to substantially reduce EED degradation, confirming the sensitivity of this system to the spatial proximity and orientation of the two ligands. As a relatively small set of potential EED degraders was evaluated in this study, ongoing efforts are aimed at determining the ‘sweet-spot’ for optimal linking within this ligand pair and establishing broader structure-degradation relationships in order to optimize the degradation efficiency of this class of molecules.

Although UNC6852 contains a potent and selective ligand for EED to mediate EED degradation, we were pleased to find that EZH2 was potently degraded in a parallel fashion, and SUZ12 to a somewhat lesser extent in multiple cell lines. This phenomenon of a bivalent degrader not only degrading its intended target, but an entire protein complex is quite unique. This result was confirmed by both western blot experiments as well as more extensive global proteomics studies, which also revealed the exquisite selectivity of UNC6852 mediated degradation within the proteome. It has been known for some time that EZH2 is not catalytically active in isolation, and recent structural studies have revealed that EZH2, EED, and SUZ12 associate intimately and the interactions between these three subunits seem to closely regulate enzymatic activity (Jiao *et al*., 2015). Specifically, EED is engulfed by a belt-like structure of EZH2, and SUZ12 contacts both of these two subunits. As a result, EZH2 is positioned in very close proximity to the EED226 binding site. Mechanistically, it is possible that UNC6852 mediates the direct ubiquitylation of EED, EZH2, and SUZ12. Alternatively, ubiquitylation of one of the three PRC2 components may result in the entire complex being recruited to the proteasome for degradation due to the close association and intertwined nature of the three proteins. It is possible that some combination of these two mechanisms contribute to overall PRC2 degradation, as EED, EZH2, and SUZ12 are not all degraded to the same extent under identical conditions. We observed that SUZ12, which is somewhat further from the EED226 binding site in PRC2 is degraded to a lesser extent than both EED and EZH2, suggesting that PRC2 is not consistently recruited to the proteasome as a single unit. Overall, these mechanistic questions are challenging to tease apart but they are of high interest conceptualizing the degradation of protein complexes more broadly.

Due to the genetic data linking PRC2 to tumorigenesis, extensive efforts have led to the development of numerous clinical candidates that target the SET domain of EZH2, as well as more recently the WD40 domain of EED. However, it has been reported that resistance to SAM-competitive EZH2 inhibitors can be caused by single point mutations in cell culture (Baker *et al*., 2015; Gibaja *et al*., 2016), suggesting that patients may become refractory to this class of molecules. Targeted protein degradation as a therapeutic approach is unique in that it is more likely to prevent the evolution of target-directed resistance mechanisms, and recent excitement over this approach to drug discovery cannot be overstated. As a result, we were motivated to investigate small molecule induced PRC2 degradation as an additional approach to targeting PRC2, particularly in the context of human cancer cell lines that are sensitive to EZH2 and EED inhibition. We demonstrate that UNC6852 has comparable antiproliferative effects to EZH2 and EED inhibitors (UNC1999 and EED226, respectively) in DB cells. We can attribute the effect observed with UNC6852 to PRC2 catalytic degradation versus on target inhibition because the negative control UNC7043 which potently binds EED *in vitro* but does not engage VHL has no effect. Importantly, the cell toxicity observed during this study with UNC6852 was substantially less than with both EED and EZH2 inhibitors, further supporting the notion that PRC2 degraders may have specific advantages over existing inhibitors. In summary, the results presented in this study demonstrate that PRC2 targeted degradation can be achieved and is a viable approach to selectively and potently inhibit PRC2 function. UNC6852 represents a useful tool compound to further interrogate PRC2 function in development and disease, as well as for further development into potential therapeutics.

## SIGNIFICANCE

The misregulation of PRC2 due to EZH2 overexpression or EZH2 gain-of-function mutations is prevalent in oncogenesis. Despite the growing number of EZH2 inhibitors in the clinic, inhibitor resistance through subsequent EZH2 mutations and chemoresistance in cancer is still a concern and new therapeutic approaches are clearly needed. We report the first example of PRC2 degradation with a bivalent chemical degrader (UNC6852). Using an EED-targeted degrader, we demonstrate the successful degradation of all core PRC2 components including EED, EZH2, and SUZ12. PRC2 degradation leads to a loss in PRC2 catalytic activity, a decrease in H3K27me3 levels, and anti-proliferative effects in a DB cell line with an EZH2^Y641N^ gain-of-function mutation. Importantly, the antiproliferative effects of UNC6852 are comparable to those of potent inhibitors of EZH2 and EED. UNC6852 provides a unique tool for studying PRC2 function and downregulation of PRC2 activity in cancer. Additionally, PRC2-targeted degraders may have the ability to overcome acquired resistance to EZH2 small molecule inhibitors and provide a complementary therapeutic strategy to compounds currently in clinical development.

## ACKNOWLEDGEMENTS

This work was supported by the National Institute on Drug Abuse, National Institutes of Health (NIH) (Grant R61DA047023-01) and the University Cancer Research Fund, University of North Carolina at Chapel Hill to L.I.J. and the National Institute of General Medical Sciences, NIH (Grant R01GM100919) to S.V.F. This research is based in part upon work conducted using the UNC Proteomics Core Facility, which is supported in part by P30CA016086 Cancer Center Core Support Grant to the UNC Lineberger Comprehensive Cancer Center. The authors thank Stephen V. Frye, Brian D. Strahl, Raghuvar Dronamraju, and Edward P. Browne for their helpful discussions throughout the project. The authors thank Ronan P. Hanley for reviewing the primary chemistry data and Jarod M. Waybright for reviewing the primary biology data. The authors thank Cristin M. Galardi for her guidance using the Jess (ProteinSimple) and Brian P. Hardy for assembly of the screening plate for TR-FRET.

## AUTHOR CONTRIBUTIONS

Conceptualization, F.P and L.I.J.; Formal Analysis, F.P., A-M.W.T., J.B., J.M.R., and L.E.H.; Investigation, F.P., A-M.W.T., J.B., J.M.R., and L.E.H; Resources, S.H.C. and J.L.N-D.; Writing – Original Draft, F.P and L.I.J.; Writing – Review & Editing, F.P. and L.I.J.; Visualization, F.P., A-M.W.T., and L.E.H.; Supervision, L.E.H., K.H.P, D.M.M., and L.I.J; Project Administration, L.I.J.; Funding Acquisition, L.I.J.

## ONLINE METHODS

### Protein Expression and Purification

EED recombinant protein (residues 1-441, accession number: AAD08714) was expressed and purified with an N-terminal His tag as previously reported (Barnash *et al*., 2017).

### Time Resolved-Fluorescence Energy Transfer Assay

The TR-FRET assay was developed and performed as previously reported (Rectenwald *et al*., 2019). Briefly, assays were run using white, low-volume, flat-bottom, nonbinding, 384-well microplates (Greiner, 784904) containing a total assay volume of 10 μL per well. The assay buffer was composed of 20 mM Tris [pH 7.5], 150 mM NaCl, 0.05% Tween 20, and 2 mM DTT. LANCE Europium (Eu)-W1024 Streptavidin conjugate (2 nM) and LANCE *Ultra* U*Light*™-anti-6x-His antibody (10 nM) were used as donor and acceptor fluorophores associated with the tracer ligand and protein, respectively. Final assay concentrations of 15 nM 6X histidine tagged EED protein (residues 1-441, N-terminal tag) and 15 nM of UNC5114-biotin tracer ligand were used for final compound testing. Assay performance was evaluated using the Z’ factor calculation at varying DMSO concentrations up to 3%. Low signals were obtained using 50 µM EED226 to obtain complete inhibition and high signals were obtained without compound. The Z’ factor was consistent at each DMSO concentration revealing a DMSO tolerance of up to 3% (Z’ 0.5% = 0.83, Z’ 3% = 0.80).

A 10 point, three-fold serial dilution of each compound at 100X final assay concentration was made in DMSO using a TECAN Freedom EVO liquid handling workstation to create an assay mother plate. The top concentration of each compound in the mother plate was 1 mM. Using a TTP Labtech Mosquito® HTS liquid handling instrument, assay ready plates were stamped with 100 nL of the compound solutions from the mother plate. 10 µL of a mixture consisting of EED, UNC5114-biotin, and the fluorophore reagents (concentrations noted above) was added to each well of an assay ready plate using a Multidrop Combi (ThermoFisher). After addition of assay components, plates were sealed with clear covers, mixed gently on a tabletop shaker for 1 minute, centrifuged at 1000xg for 2 minutes, and allowed to equilibrate in a dark space for 1 hour. After 1 hour, the plate was read on an EnVision 2103 Multilabel Plate Reader (PerkinElmer) using an excitation filter at 320 nm and emission filters at 615 and 665 nm. Emission signals (615 and 665 nm) were measured simultaneously using a dual mirror D400/D630 (using a 100-microsecond delay). TR-FRET output signal was expressed as emission ratios of acceptor/donor (665/615 nm) counts. Percent inhibition was calculated on a scale of 0% (i.e., activity with DMSO vehicle only) to 100% (100 μM EED226) using full column controls on each plate. The data was fit with a four-parameter nonlinear regression analysis using GraphPad Prism to determine IC_50_ values and are reported as an average of two biological replicates ± standard deviation.

### Cell Culture and Lysis

Cells were maintained in a humidified incubator at 37°C, 5% CO_2_. HeLa cells (ATCC) were cultured in MEM-α 1X (Gibco™, 12571071), 10% FBS (VWR Seradigm, 89510-194), and 1% non-essential amino acids (Gibco™, 11140050). DB cells (ATCC, CRL-2289™) were cultured in RPMI 1640 (Gibco™, 11-875-093). 293T cells (ATCC, CRL-11268™) were cultured in DMEM (Gibco, 11995-065), 1% pen/strep, and 10% FBS. For degradation analysis, cells were cultured in 6 well plates (Olympus Genesee Scientific, 25-105) and dosed with the appropriate concentration of bivalent degrader from a DMSO stock. Adherent cells (HeLa) were seeded at 400,000 cells/well for 24 hr analysis and 100,000 cells/well for 72 hr analysis. At the appropriate time point, cells were washed with 2X PBS, scraped in PBS (1 mL), centrifuged, aspirated, and lysed in 40-50 μL of modified RIPA lysis buffer (1X Modified RIPA buffer (25 mM Tris pH 8, 150 mM NaCl, 1 % NP-40, 1 % sodium deoxycholate, 0.1 % SDS), 1X Protease inhibitor cocktail (Active Motif, 37490), 4μL/mL Benzonase® nuclease (Millipore, ≥90% SDS page, E1014), DPBS (Gibco™)). Non-adherent cells (DB) were seeded at 800,000 cells/well for 24 hr analysis, and 100,000 cells/well for 72 hr analysis. Cells were centrifuged, aspirated, washed with 2X PBS, and aspirated again and lysed in 40-50 μL of Cytobuster lysis buffer (Cytobuster™ (71009), 1X Protease inhibitor cocktail (Active Motif, 37490), 2μL/mL Benzonase® nuclease (Millipore, ≥90% SDS page, E1014)).

The protein levels were quantified using Pierce™ Detergent Compatible Bradford Assay Kit (Thermo Scientific, 23246) for the modified RIPA lysis buffer, and with Protein Assay Dye Reagent Concentrate (Bio-Rad, 5000006) using a known concentration of BSA standard for the Cytobuster lysis buffer.

### Jess ProteinSimple Analysis

Jess Protein Simple was used according to product guideline instructions. HeLa cell were treated with bivalent degraders UNC6851, UNC6852, UNC6853, UNC6845, UNC6846, UNC6847 at 5 μM for 4, 24, and 48 hrs, and cell lysates were generated using a modified RIPA buffer. Cell lysates were used at 1mg/mL. The primary antibodies used were: anti-EED (1:10, R&D Systems, AF5827), anti-EZH2 (1:100, D2C9 XP®, Cell Signaling Technology, 5246S). The secondary antibodies used were: anti-sheep HRP secondary antibody (1:25, LifeTech/Novex, A16041), anti-rabbit secondary IR antibody (1:20, Protein Simple, 043-820).

### Western Blot Analysis

Cell lysate (20 μg) was combined with Laemmli buffer (Bio-Rad; 2X – 1610737 or 4X – 1610747) containing 2-mercaptoethanol (5%) and samples were boiled at 95 °C prior to gel loading. Gels (15μL, 15 well; 4-15% precast mini-PROTEAN® TGX™ gels, Bio-Rad, 4561046DC; or 4-15% precast mini-PROTEAN® TGX™ Stain Free™ gels, Bio-Rad 4568086) were placed in a Mini-PROTEAN® tetra cell at 200V in 1X Tris/Gycine/SDS running buffer (Bio-Rad, 1610772). Molecular weight ladder’s used were either Precision Plus Protein™ Dual Color Standard (Bio-Rad, 161-0374), or PageRuler™ Plus pre-stained protein ladder (ThermoFisher, 26619). Protein was transferred onto Immobilon-FL PVDF Membranes (Millipore Sigma, IPFL00010), with 1X Tris/Gycine transfer buffer (Bio-Rad, 1610772) and methanol (0.2% volume) at 100V for 1 hr at 4°C. Membranes were blocked at room temperature for 1 hr with Odyssey® blocking buffer (TBS, LI-COR, 926-31099), and the incubated with primary antibodies overnight at 4°C.

#### Primary Antibodies

anti-EED (1:500, R&D Systems, AF5827), anti-EZH2 (1:1000, D2C9 XP® Cell Signaling Technology, 5246S), anti-SUZ12 (1:500, D39F6 XP® Cell Signaling Technology, 3737S), anti-GAPDH-AlexaFluor® 680 (1:5000, Abcam, ab184095), anti-GAPDH (1:5000, EMD Millipore, AB2302), anti-H3K27me3 (1:2000, Abcam, ab6002), anti-Histone H3 (1:5000, Abcam, ab1791), anti-HA (Abcam, 1:500, ab9110).

Membranes were incubated with the corresponding secondary antibodies for 1hr at room temperature prior to imaging. Fluorescence imaging was performed on a LI-COR Odyssey. For chemiluminescent detection, membranes were activated with ECL Prime western blotting detection reagent (Amersham, RPN2232) and imaged on a Bio-Rad Chemidoc.

#### Li-COR Fluorescent Secondary Antibodies

IR Dye® 680RD (1:10000, Goat anti-mouse, LI-COR, 926-68070), IR Dye® 800CW (1:10000, Goat anti-rabbit, LI-COR, 926-32211).

#### Chemi-Doc Chemiluminescent Secondary Antibodies

Goat anti-chicken HRP (1:10000, LifeTech/Novex, A16054), Donkey anti-sheep HRP (1:10000, LifeTech/Novex, A16041), Donkey anti-rabbit HRP (1:10000, LifeTech/Novex, A16035).

### Western blot quantification

Western blots were analysed by firstly calculating the densitometry on either ImageStudio software or ImageLab software for LI-COR or Chemidoc imaging, respectively. The densitometry of the protein of interest band relative to the densitometry of each corresponding GAPDH band was calculated. The resulting densitometry relative to the DMSO band was calculated to give the % degradation.

For the dose response and time study these values were plotted in GraphPad Prism against the corresponding concentration or time of degrader treatment. An inhibitor concentration vs response (three parameters) regression was plotted and the IC_50_ values were taken from GraphPad Prism which corresponded to either the apparent half-life (t_1/2_, when protein levels were plotted against time), or the half maximal degradation concentration (DC_50_, when protein levels were plotted against concentration). The maximal degradation (D_max_) was calculated based on the % degradation at 30 μM after 24 hours).

### Cell Proliferation Analysis

Exponentially growing DB cells were seeded in a 12 well plate (Corning® Costar®, CLS3513) at a cell density of 0.5 × 10^5^ cells/mL. Every 3 days the media was exchanged, cells were split back to the seeding density, and the compound or DMSO control were re-dosed. At each time point the cells were counted on an automated Bio-Rad TC20™ cell counter with Trypan blue (Abcam, ab233465) and cell counting slides (1450015) to give the cell count (cells/mL) and cell viability (%). The % cell proliferation is calculated based on the total cell number expressed as split-adjusted viable cells, relative to the DMSO control at the same time point. To determine an EC_50_, the total cell number is expressed as a split-adjusted viable cells/mL and the results were analyzed in GraphPad Prism with a log(inhibitor) vs response – variable slope (four parameters). Experiments were performed in biological triplicate.

### Antibody Validation Methods

pCMVHA EED WT (Addgene plasmid # 24231; http://n2t.net/addgene:24231; RRID:Addgene_24231) and pCMVHA hEZH2 (Addgene plasmid # 24230; http://n2t.net/addgene:24230; RRID:Addgene_24230) were a gift from Kristian Helin.(Bracken *et al*., 2003) Constructs were confirmed by sequencing prior to use. pCMVHA EED WT contains the DNA sequence to express the two smaller isoforms of EED (EED3/4) but not the two larger predicted isoforms EED1/2.(Montgomery *et al*., 2007) Two individually isolated plasmids from the original bacterial streak received from Addgene were transfected into 293T cells using Fugene HD (Promega) per manufacturer’s protocol. Cells were collected two days post transfection and assayed by western blot using anti-HA (Abcam, ab9110, 1:1500) to confirm protein expression. For validation of EED antibody AF5827 (R&D Systems) and EZH2 antibody CST5246 (Cell Signaling Technology), the same plasmids were transfected into HeLa cells using Fugene HD and cell lysates analyzed by western blot using the respective antibodies relative to untransfected controls. Blots are provided overexposed to see endogenous levels of EED and EZH2 in untransfected controls as well as overlaid on membrane to allow for comparison to the ladder (PageRuler Plus Prestained Protein Ladder, 10-250kDa, ThermoFisher). GAPDH (Millipore, AB2302, 1:5000) is provided as a loading control for all samples.

### Global Proteomics Experiments

Exponentially growing HeLa cells were seeded in 10 cm plates and treated with UNC6852, UNC7043, or DMSO (10 μM, 24 hrs). Cells from biological triplicates were harvested on ice and centrifuged at 4 °C for 10 min. Lysis on ice with 8M urea in 50 mM Tris-HCl pH 7.8 with 1X protease inhibitor cocktail (Active Motif, 37490) and 1X phosphatase inhibitor (Roche, PhosSTOP, 05892791001). Sample preparation: Each sample (200 μg) was incubated with trypsin overnight. Samples were desalted with SepPak C18 cartridges (Waters, 100 mg sorbent, WAT036820), and Pierce BCA peptide quantitation assay was performed. Each sample (50 μg) was labelled with a TMT label. After labelling, samples were quenched and combined 1:1 into a single multiplexed sample. An aliquot (100 μg) of the mixed sample was fractionated into 8 fractions using the Pierce high pH reversed phase fractionation spin columns. Peptide fractions were analysed in duplicate by LC-MS/MS using a Thermo Easy nLC 1200-QExactive HF. Proteins were identified by searching raw data against a reviewed Uniprot human database (containing 20,245 sequences) using Andromeda and quantified using TMT intensities within MaxQuant v1.6.3.4. Further data analysis was performed in Perseus, Excel, and GraphPad Prism. Statistical significance between each pair of groups was calculated using Student’s T-test and a p-value of <0.01 was used as the significance cut-off. Log2 fold change of each protein was calculated by dividing the averaged log2 TMT intensities of each compound by the averaged log2 TMT intensities of the DMSO control across all replicates. A log2 absolute fold change of 0.5 was used as the significance cut-off.

## SUPPLEMENTAL FIGURES

**Figure S1.**
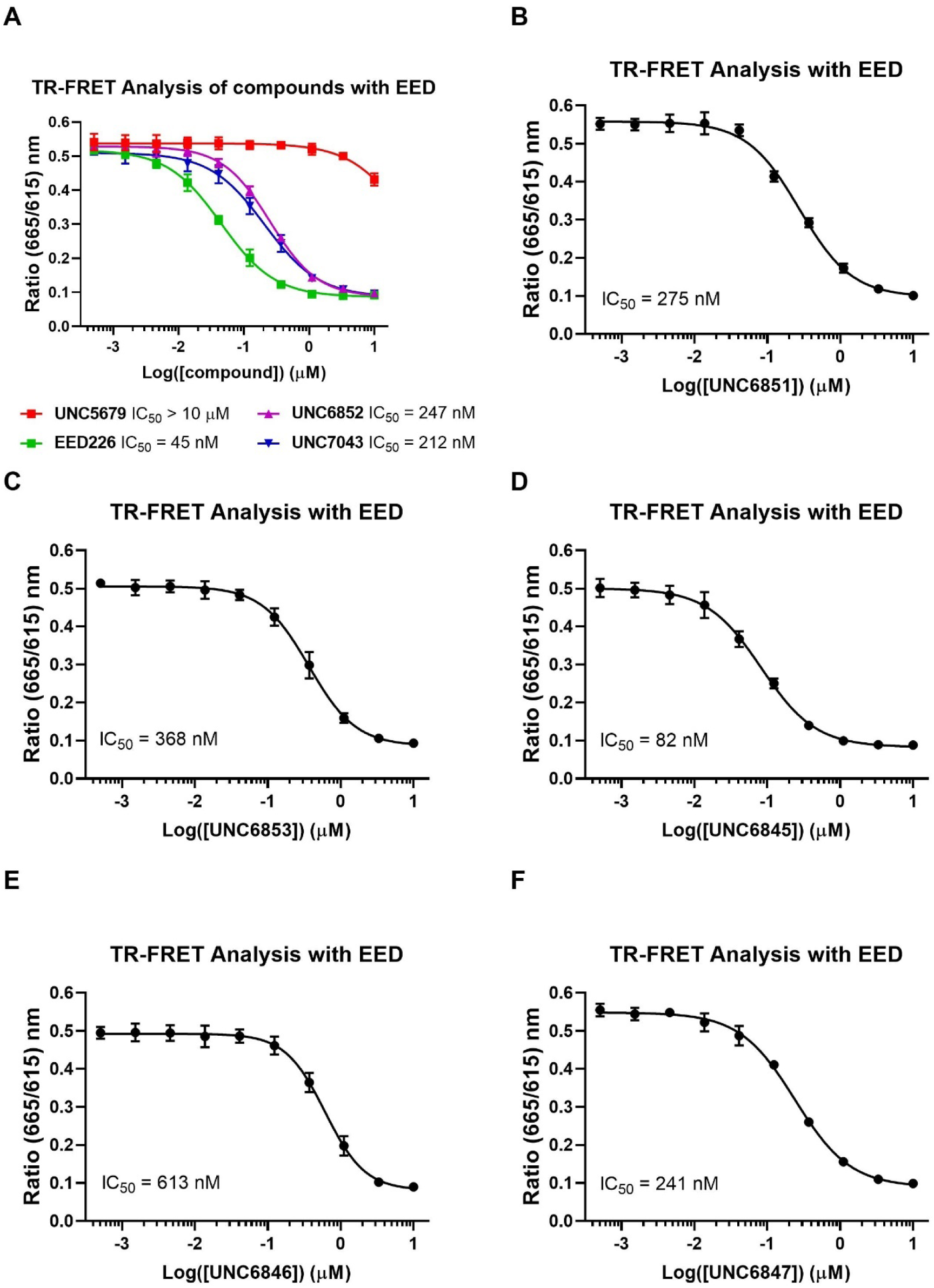
TR-FRET dose response curves of bivalent degraders binding to recombinant EED (related to Table 1). IC_50_ values were determined by TR-FRET and are reported as the average of two biological replicates ± standard deviation. (**A**) EED226 (IC_50_ = 45 ± 6.9 nM), UNC6852 (IC_50_ = 247 ± 2.90 nM), UNC7043 (IC_50_ = 212 ± 30.8 nM), UNC5679 (EED small molecule negative control, IC_50_ > 10 μM). (**B**) UNC6851 (IC_50_ = 275 ± 2.55 nM). (**C**) UNC6853 (IC_50_ = 368 ± 56.0 nM). (**D**) UNC6845 (IC_50_ = 82 ± 3.8 nM). (**E**) UNC6846 (IC_50_ = 613 ± 73.8 nM). (**F**) UNC6847 (IC_50_ = 241 ± 0.778 nM).

**Figure S2.**
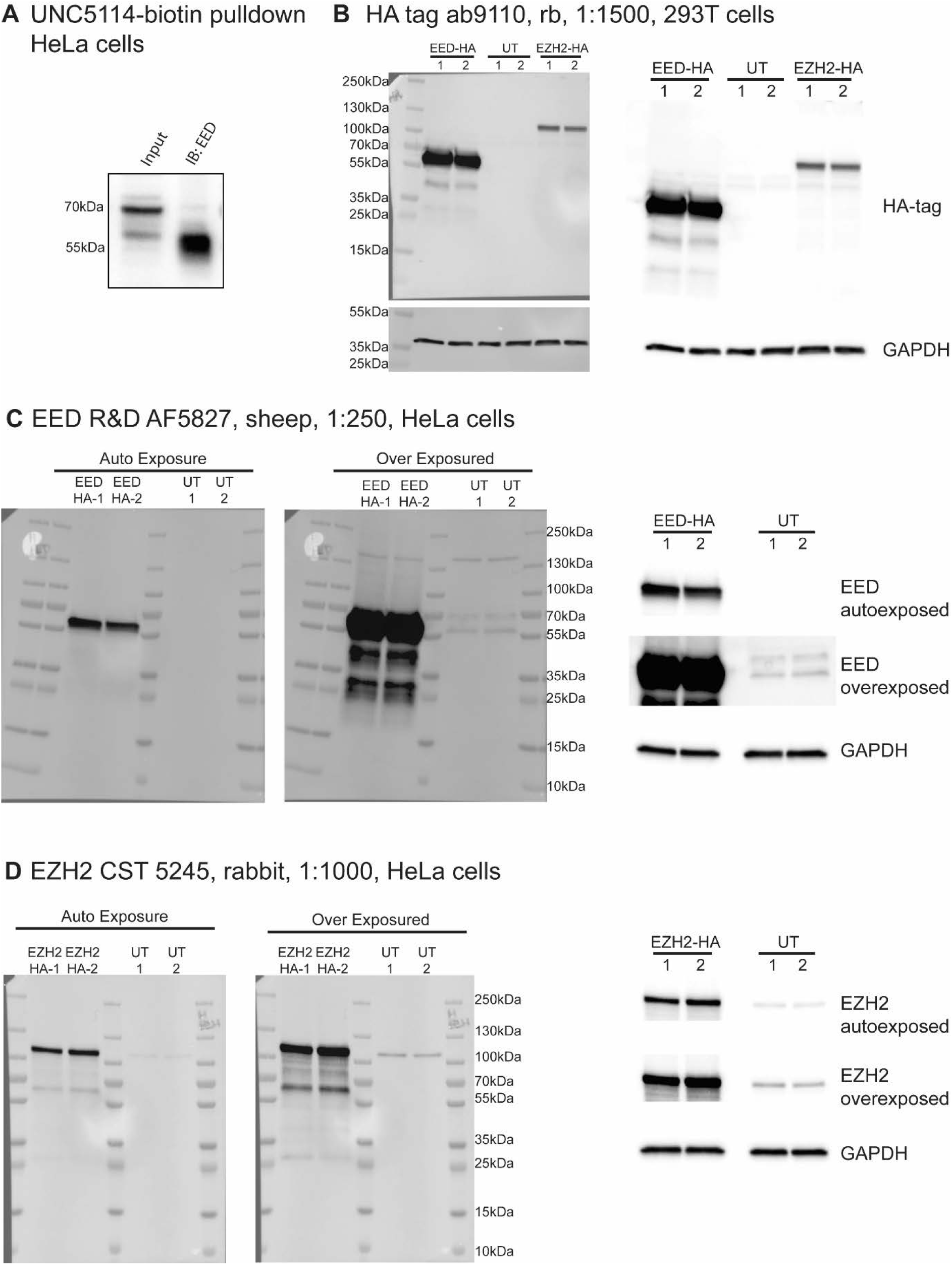
EED and EZH2 antibody validation (related to Figure 2). (**A**) Chemiprecipitation of endogenous EED protein from HeLa cell lysates with UNC5114-biotin (previously reported biotinylated EED ligand) (Barnash *et al*., 2017). Immunoblotting for EED with anti-EED (1:500, R&D Systems, AF5827) indicates that the lower band is EED. (**B**) 293T cells were transfected with HA-tagged overexpression constructs for EED and EZH2 in duplicate and lysates were analyzed for protein expression using an anti-HA antibody. Lysates from untransfected (UT) cells were run in parallel. GAPDH is provided as a loading control. (**C**) HeLa cells were transfected with HA-tagged EED in duplicate and lysates were analyzed for protein expression using anti-EED AF5827 from R&D Systems. AF5827 recognizes the HA overexpression construct and correlates to the lower band observed at ∼60 kDa in **A** and the untransfected controls, validating this band as EED, most likely isoforms 3 and 4. The overexpression construct did not contain the DNA sequence necessary to express EED isoforms 1 and 2. (**D**) HeLa cells were transfected with HA-tagged EZH2 in duplicate and lysates were analyzed for protein expression using anti-EZH2 CST5246 from Cell Signaling Technology. CST5246 recognizes the HA overexpression construct at the same molecular weight as endogenous EZH2 in the untransfected controls.

**Figure S3.**
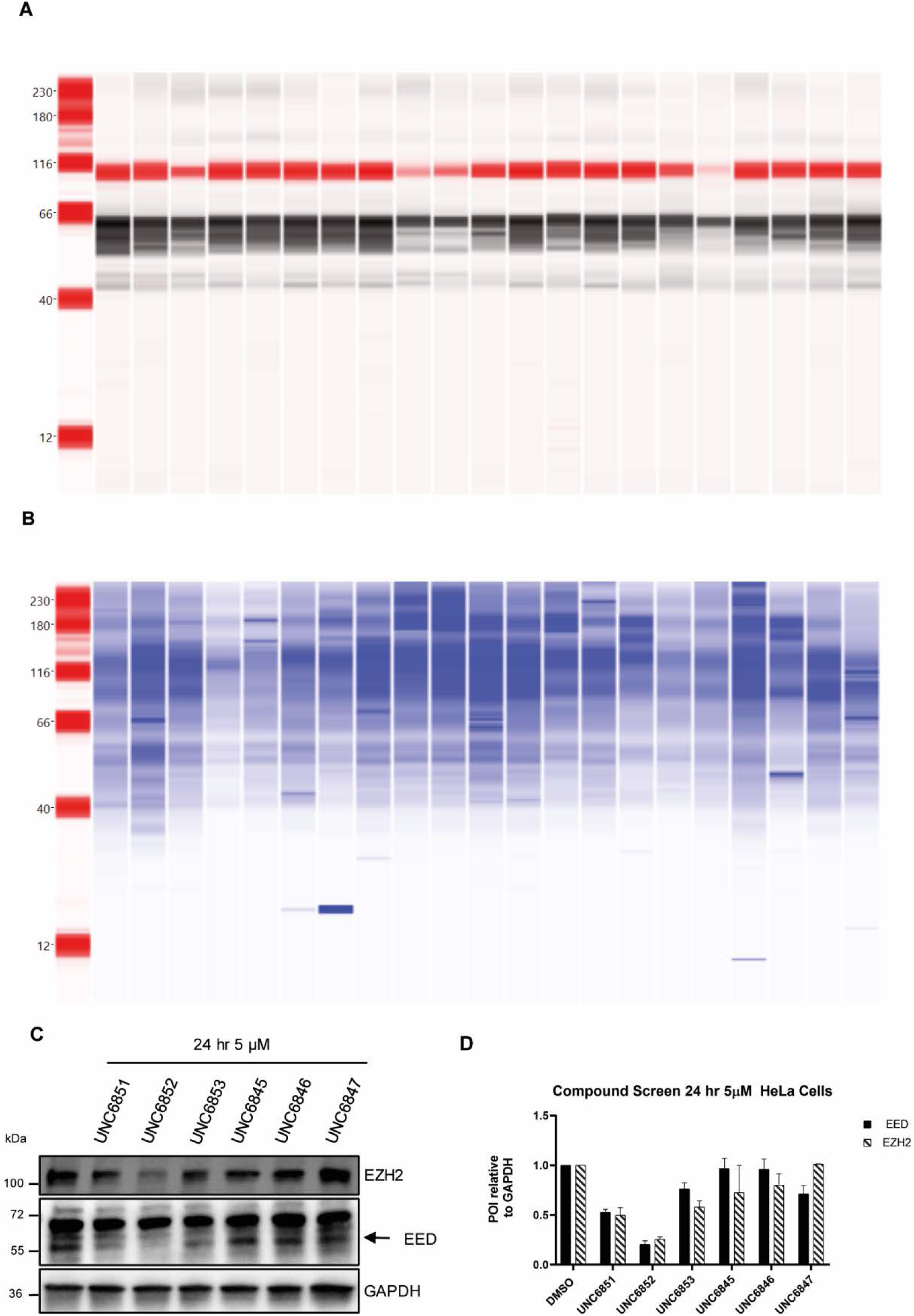
Jess primary degradation screen, western blot analysis, and quantification of EED and EZH2 degradation with 6 EED-targeted bivalent degraders (related to Table 1). (**A**) HeLa cells were dosed at 5 μM for 4, 24, and 48 hr and then lysed and run on the Jess Protein Simple (1 mg/mL). Proteins analyzed were EZH2 (Rb, IR channel, red) and EED (Sp, chemiluminescent channel, black). (**B**) Protein Normalization for results in **A**. **1** – DMSO 4 hrs; **2** – UNC6851 5 μM 4 hrs; **3** – UNC6852 5 μM 4 hrs; **4** – UNC6853 5 μM 4 hrs; **5** – UNC6845 5 μM 4 hrs; **6** – UNC6846 5 μM 4 hrs; **7** – UNC6847 5 μM 4 hrs; **8**- DMSO 24 hrs; **9** – UNC6851 5 μM 24 hrs; **10** – UNC6852 5 μM 24 hrs; **11** – UNC6853 5 μM 24 hrs; **12** – UNC6845 5 μM 24 hrs; **13** – UNC6846 5 μM 24 hrs; **14** – UNC6847 5 μM 24 hrs; **15** – DMSO 48 hrs; **16** – UNC6851 5 μM 48 hrs; **17** – UNC6852 5 μM 48 hrs; **18** – UNC6853 5 μM 48 hrs; **19** – UNC6845 5 μM 48 hrs; **20** – UNC6846 5 μM 48 hrs; **21** – UNC6847 5 μM 48 hrs. (**C**) Western blot analysis of EED and EZH2 levels following treatment of HeLa cells with 6 bivalent degraders (UNC6851, UNC6852, UNC6853, UNC6845, UNC6846, and UNC6847, 5 µM) for 24 hours. (**D**) Quantification of EED and EZH2 protein levels in **C** relative to GAPDH and plotted relative to the DMSO control. Densitometry analysis was performed with Image Lab and Image Studio. Values are the average of two biological replicates ± standard deviation.

**Figure S4.**
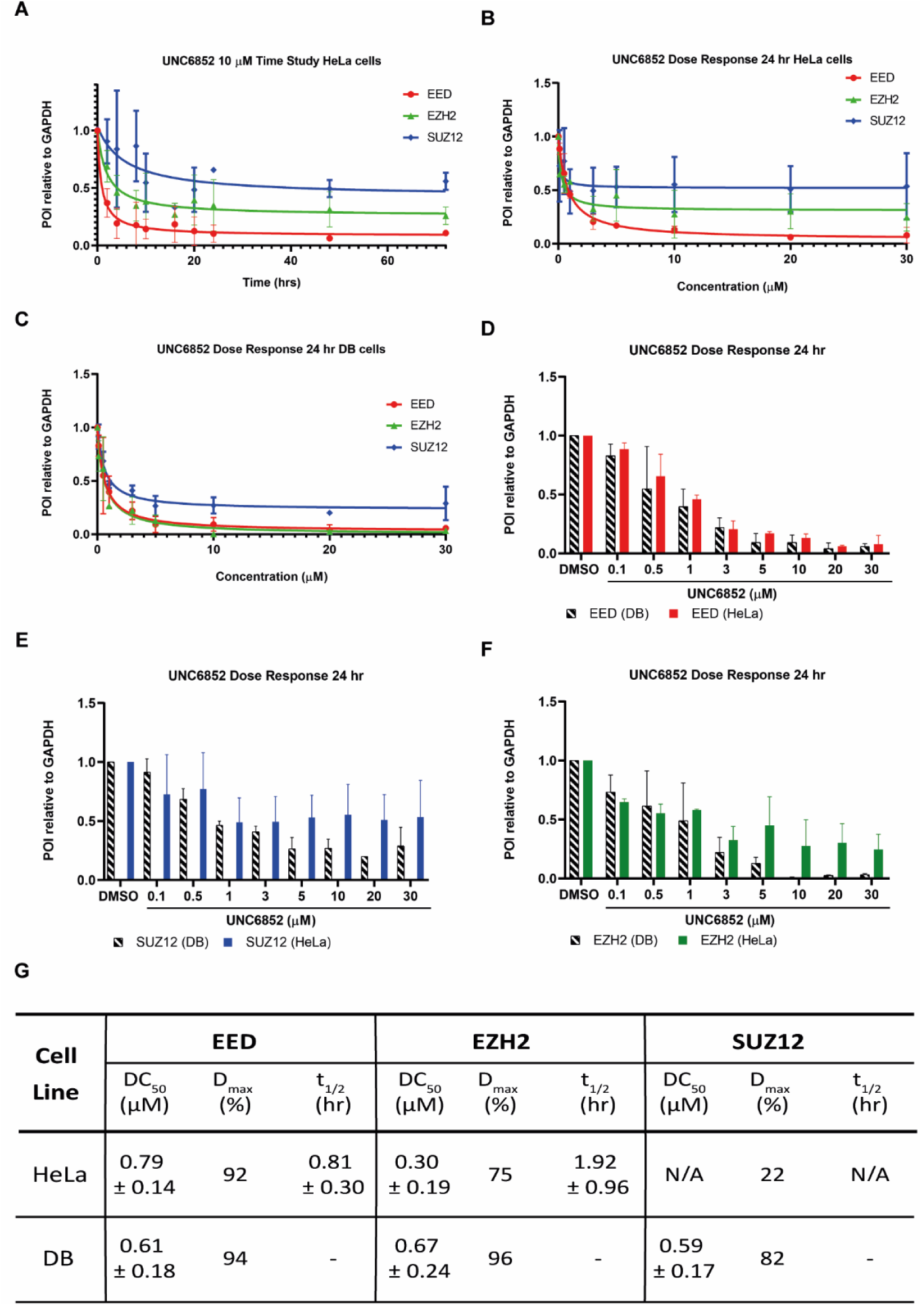
Quantification of EED, EZH2, and SUZ12 degradation and calculation of DC_50_, D_max_, and t_1/2_ in HeLa and DB cell lines (related to Figures 2 and 5). (**A**) Quantification of PRC2 degradation over time upon treatment with UNC6852 in HeLa cells (2 – 72 hrs). (**B**) Quantification of PRC2 degradation upon treatment with UNC6852 in HeLa cells for 24 hours in a dose response fashion (0 – 30 μM) (Figure 2). (**C**) Quantification of PRC2 degradation upon treatment with UNC6852 in DB cells for 24 hours in a dose response fashion (0 – 30 μM) (Figure 2). (**D**) Comparison of EED protein levels as reported in **B** (HeLa cells) versus **C** (DB cells). (**E**) Comparison of SUZ12 protein levels as reported in **B** (HeLa cells) versus **C** (DB cells). (**F**) Comparison of EZH2 protein levels as reported in **B** (HeLa cells) versus **C** (DB cells). (**G**) DC_50_ and t_1/2_ values were calculated for each protein by plotting the protein densitometry by western blot analysis relative to GAPDH, and then relative to the DMSO control, against either time (apparent half-life, t_1/2_) or concentration (DC_50_). Data is the average of two biological replicates ± standard error.

**Figure S5.**
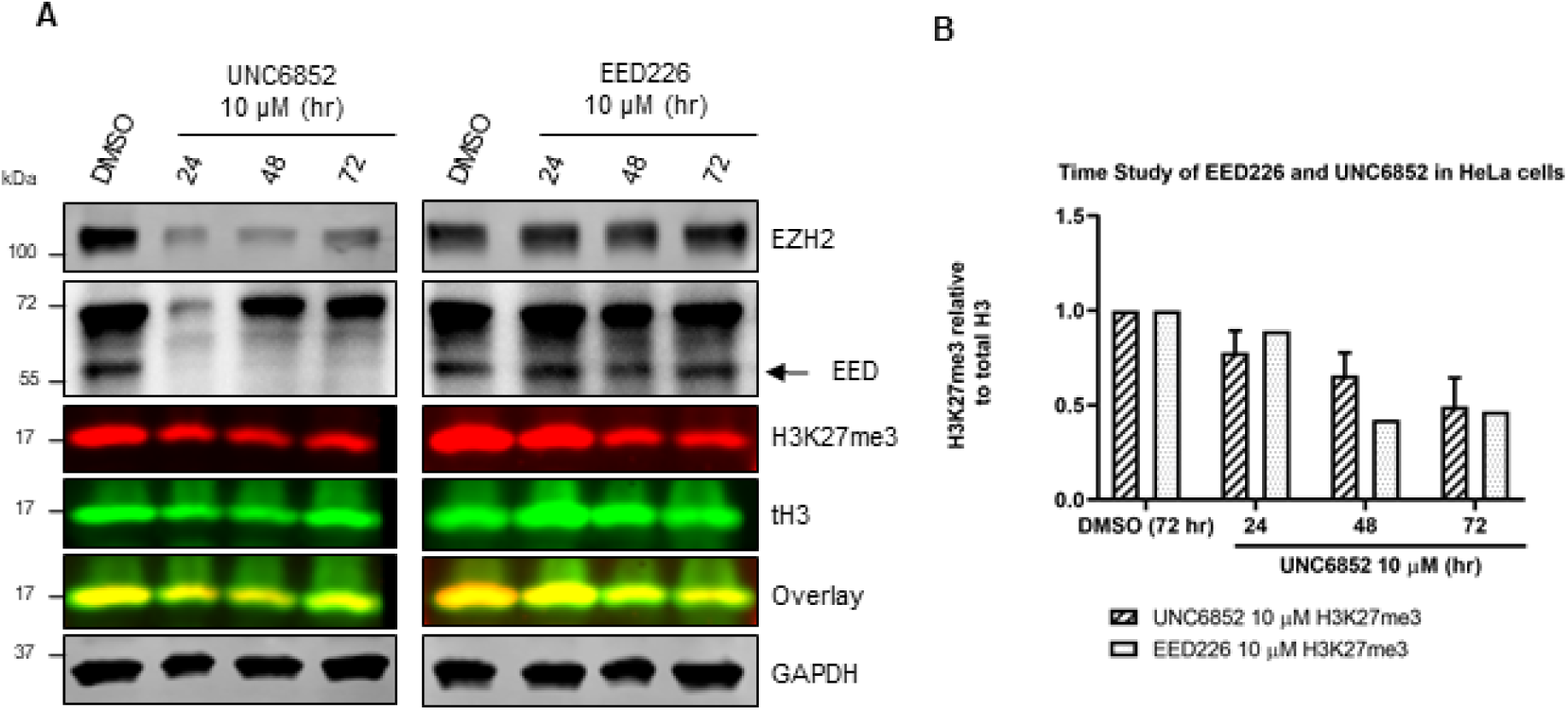
Western blot analysis and quantification of H3K27me3 levels upon treatment of HeLa cells with UNC6852 and EED226 (related to Figure 5). (**A**) Western blot analysis of EZH2, EED, and H3K27me3 levels following treatment of HeLa cells with UNC6852 and EED226 (24, 48, and 72 hrs, 10 μM). (**B**) Quantification of H3K27me3 levels in **A**. H3K27me3 levels are reported relative to total H3 and plotted relative to the DMSO control.

**Figure S6.**
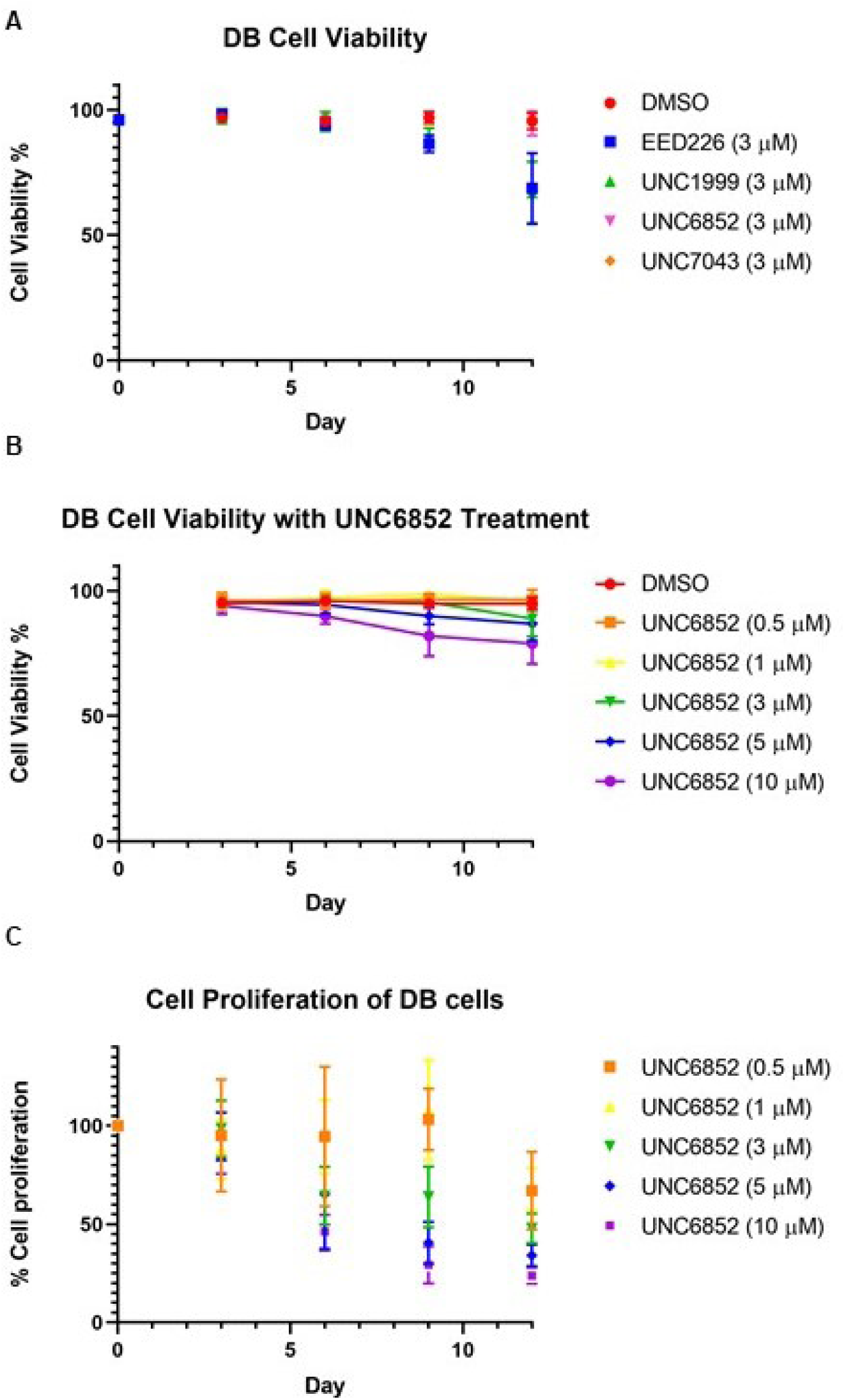
Cell viability and cell proliferation in DB cells (related to Figure 5). (**A**) Cell viability over 12 days in DB cells corresponding to Figure 5E. Cells were treated with DMSO, EED226, UNC1999, UNC6852, or UNC7043 (3 μM). Viability was evaluated with a TC20 Bio-Rad cell counter using a trypan blue stain. (**B**) Cell viability over 12 days in a DB cell line corresponding to Figure 5F. Cells were treated with UNC68522 (0.5 – 10 μM) for 12 days. Viability calculated as in **A**. (**C**) UNC6852 displays a concentration dependent inhibition of DB cell proliferation after 12 days of treatment (0.5 – 10 μM).

